# cTBS Over the Left Supplementary Motor Area (lSMA) Does Not Modulate Rhythmic Bimanual Coordination in the Presence and Absence of Visual Cues

**DOI:** 10.1101/2023.06.10.544481

**Authors:** Jaskanwaljeet Kaur, Ramesh Balasubramaniam

## Abstract

Bimanual coordination modes, namely in-phase and anti-phase, represent two distinct movement patterns characterized by simultaneous & symmetrical movements of both hands and alternating complementary actions, respectively. These coordination modes are integral in various activities, such as playing musical instruments, typing, and participating in sports that demand precise hand-eye coordination. The objective of the present experiment was to investigate the impact of continuous theta burst stimulation (cTBS) targeting the left supplementary motor area (lSMA) on bimanual coordination during in/anti-phase coordination modes. To explore this, we utilized a steady-state system of coordination dynamics and evaluated the continuous relative phase (ϕ) and variability of relative phase (SDϕ) during cued and non-cued trials in both pre- and post-transcranial magnetic stimulation (TMS) conditions. The results revealed that visual cues (cued trials) significantly enhanced bimanual coordination performance in both in/anti-phase coordination modes. However, contrary to expectations, the downregulation of lSMA through cTBS did not lead to significant disruptions in movement during in/anti-phase bimanual coordination in pre- and post-TMS stimulation. Potential factors for the lack of observed effects include methodological limitations, individual differences, and functional redundancy within the motor system. Further research is needed to optimize stimulation parameters, increase sample sizes, and explore the interactions between the lSMA, and other brain regions involved in motor control to gain a comprehensive understanding of the contributions of the lSMA in bimanual coordination.

## Introduction

Bimanual coordination substantially contributes to human movement and has been characterized by precise spatial and temporal interaction between limbs (Swinnen, 2002). Rhythmic bimanual coordination between upper limbs, i.e., the left and the right hand, can be performed simultaneously (in-phase movement pattern, where the phase difference is 0^0^) or in the opposite direction (anti-phase movement patten, where the phase difference is 180^0^). It has been shown that these movement patterns are easily maintained at low cycling frequencies (Haken et al., 1985). The involvement of interhemispheric connections appears to be important for bimanual coordination and has been widely discussed (Geffen et al., 1994; Kazennikov et al., 1999; Gazzaniga, 2000; Donchin et al., 2002; Swinnen & Wenderoth, 2004; Chen et al., 2005; Aramaki et al., 2006; Nachev et al., 2008; Y. L. Kermadi E. M. Rouiller, I., 2000; Liuzzi et al., 2011; Jung et al., 2020; Miyaguchi et al., 2020). In fact, prior studies have shown the importance of interhemispheric connections via callosal contributions in bimanual key press task (Tanji et al., 1988), bimanual tapping tasks (Leonard et al., 1988; Bonzano et al., 2008) and in continuous circle drawing task (Kennerley et al., 2002).

In addition to interhemispheric connections, there seems to be further involvement of specific brain regions during coordination of bimanual movements. In particular, the supplementary motor area (SMA), located in the medial part of the premotor cortex constituting Broadmann’s area 6, has been shown to be involved in motor sequencing, spatial/temporal processing, working memory and music processing (Cona & Semenza, 2017). An fMRI study has further shown that the SMA was active when participants performed sequencing and continuous bimanual movements (Toyokura et al., 2002). Single cell recordings from monkeys have also suggested the role of SMA in control of sequential bimanual coordinated movements (I. Kermadi et al., 1998). Based on prior work, the SMA is a cortical structure that plays a significant role in motor sequencing and execution of continuous bimanual movements.

In recent years the technique of transcranial magnetic stimulation (TMS) has been utilized to non-invasively stimulate different regions of the brain, allowing researchers to causally affect neural firing in a participant’s brain to induce certain patterns of electrical activity. Causal manipulations via TMS affect firing rate of pre-synaptic neurons and can cause post-synaptic changes in corticospinal excitability (Bestmann & Krakauer, 2015). Previously, different variations of TMS, i.e., stimulation intensity, duration etc. have been employed to study the effects of stimulation on bimanual coordination. For instance, Obhi et al., 2002 and Serrien et al., 2002 have shown that repetitive TMS (rTMS) to the SMA disrupts and degrades movement during bimanual coordination. Steyvers et al., 2003 used high-frequency rTMS of the SMA and focused on the quality of coordination during cyclical bimanual movements, specifically during in-phase and anti-phase movements and showed that after rTMS, the mean relative phase error between hands increases during anti-phase trials. More recently, researchers have investigated SMA’s role in on control of bimanual coordination using electroencephalography (EEG), showing that the pre-movement cortical oscillatory coupling within the motor network, including beta-band phase synchrony in a bi-hemispheric primarily motor cortices and spectral power at SMA, influences bimanual coordination stability (Iwama et al., 2022). These findings thus suggest that by monitoring and potentially enhancing the brain’s preparatory activity, it may be possible to improve bimanual coordination in individuals with motor impairments.

The SMA has also been shown to play a crucial role in internal and external movement generation (Debaere et al., 2001; Cunnington et al., 2002; Debaere et al., 2003; Therrien et al., 2012). In terms of internal movement generation, the SMA is involved in the planning and execution of voluntary movements requiring coordination of both hands. Prior studies have shown that the SMA is particularly active during the preparation phase of bimanual movements, such as when an individual plans to perform a specific sequence of movements with both hands simultaneously (Cunnington et al., 2002; Toyokura et al., 2002). The internal generation of bimanual movement involves the integration of motor commands and the coordination of motor representations within the SMA (Thaler et al., 1988). On the other hand, the SMA has also been shown to be involved in external generation of bimanual movements, which refers to movements that are externally cued or guided by external stimuli (Thickbroom et al., 2000; Picard & Strick, 2003; Debaere et al., 2003). In this case, the SMA contributes to coordination and synchronization of bimanual movements based on external cues. For example, when individuals perform mirror movements, the SMA ensures the temporal and spatial coordination of movements (Serrien et al., 2002; Wilson et al., 2014).

Thus, based on prior literature implicating the roles of SMA in bimanual coordination tasks, we hypothesized the downregulating the left SMA (lSMA) region via cTBS would negatively impact bimanual in-phase and anti-phase movements between the left and the right arms. We focus specifically on the lSMA due to its established role in bimanual coordination tasks and its involvement in fine motor skills (Schramm et al., 2019), indicating that movement in the contralateral limb would be affected during lSMA stimulation. Previous research has also highlighted the importance of bilateral SMA function in coordination bimanual movements and facilitating interhemispheric communication during such tasks (Carson, 2005; Calvert & Carson, 2022). To test the influence of lSMA during bimanual coordination, we utilized a novel in-phase and anti-phase bimanual task developed via the Kinarm robotic exoskeleton (BKIN Technologies Ltd, Ontario, Canada). This task allowed us to investigate bimanual coordination performance in two conditions: one with visual cues (cued trials) and one without visual cues (non-cued trials).

By incorporating both conditions in the experiment, one with cued trials and one with non-cued trials, we aimed to elucidate the impact of transiently disrupting the function of the lSMA on bimanual coordination performance. We predicted that the performance of bimanual task with visual cues, representing the externally cued condition, will be less adversely affected by lSMA disruption compared to the bimanual task without visual cues, representing the non-cued condition. This hypothesis is grounded in the understanding that the SMA is essential in synchronizing bimanual movements based on external cues, such as those provided by the visual cues (cued condition) in this study (Steyvers et al., 2003). Therefore, we anticipated that disrupting the lSMA function may impair a participants’ ability to effectively utilize their internal sense of timing and spatial coordination in the absence of external cues, leading to increased difficulty in achieving accurate and synchronized bimanual movements (Thickbroom et al., 2000; Macar et al., 2004). Furthermore, we expected that the bimanual task without visual cues, which relies more on internal generation processes within the SMA, will be more significantly affected by the downregulation of the lSMA. This is because participants will rely solely on their internal sense of timing and spatial coordination without the aid of external cues, thereby making the task more challenging and potentially prone to disruptions caused by lSMA downregulation. By contrasting the continuous relative phase (ϕ) and variability of relative phase (SDϕ) differences between the baseline task performance and performance after cTBS, we aimed to gain insights into the influence of lSMA disruption on bimanual coordination.

## Methods

### Participants

Thirty-nine subjects provided their informed consent to participate in the study. Among them, data from fifteen participants was excluded due to incomplete study participation, noncompliance with task instructions or difficulty locating the motor hotspot prior to cTBS. The remaining twenty-four participants (age: 21.46 ± 2.70, 18 female) completed both sessions of the study and were included in the data analysis. The experiment was conducted in accordance with the Declaration of Helsinki and was approved by the Institutional Review Board of the University of California, Merced.

### Handedness Measurements

Handedness was assessed by having participants complete a 4-item Edinburgh Handedness Inventory (EHI) – Short Form (Veale, 2014). The EHI accesses hand dominance in daily activities (e.g., writing, throwing). The laterality quotient (LQ) of hand dominance ranges from −100 (left-handed) to 100 (right-handed): an LQ between −100 & −61, −60 & 60, and 61 and 100 were considered left handers, mixed handers, and right handers, respectively. In the present study all the subjects were classified as right-handed.

### Experimental Design

In the present study, we used the Kinarm upper limb robotic exoskeleton. The Kinarm device is a specialized apparatus that facilitates the assessment of upper limb function. It’s composed of a height-adjustable chair that has bilateral arm and hand support platforms. The device also includes a monitor connected to the operator’s computer and a display screen that is located underneath the monitor. This setup allows participants to perform two-dimensional movements while simultaneously observing and interacting with the visual stimuli projected onto the screen. The Kinarm device enables the assessment of both visual stimuli and bimanual arm movements within the same workspace. During the task, the Dexterit-E software continuously records the participants’ movements at a sampling rate of 1000hz (3.8v, BKIN Technologies Ltd, Ontario, Canada). The software captures the hand position coordinates (x, y) as well as the movement velocity and acceleration of the arm along the transverse plane. Upon completion of the experiment, the recorded data is automatically saved as a c3d data file.

### In-phase & Anti-phase Coordination Modes

We employed a previously developed bimanual coordination task (Kaur et al., 2023), implemented using Simulink with a few modifications (R2015a, The MathWorks, USA), to investigate the effects of TMS on continuous in-phase and anti-phase coordination during pre- and post-TMS stimulation. The start position of the task is illustrated in Figure 1b, with four target circles positioned 5cm apart on the screen in front of the participant. The experimental setup for both coordination modes (in-phase and anti-phase) was similar, but the movements differed depending on the coordination mode being tested. Prior to beginning the task, participants were trained on the coordination mode they were randomly assigned to, either in-phase or anti-phase. Participants moved their hands to the white targets (initial starting position), prompting the red targets to flash five times, signaling the start of a new trial. The cycling frequency, also known as the speed of movement, was set at 750ms, which was determined based on piloting the experiment with adult (>18 years) participants, as it was considered a comfortable speed of movement.

**Figure 1.**
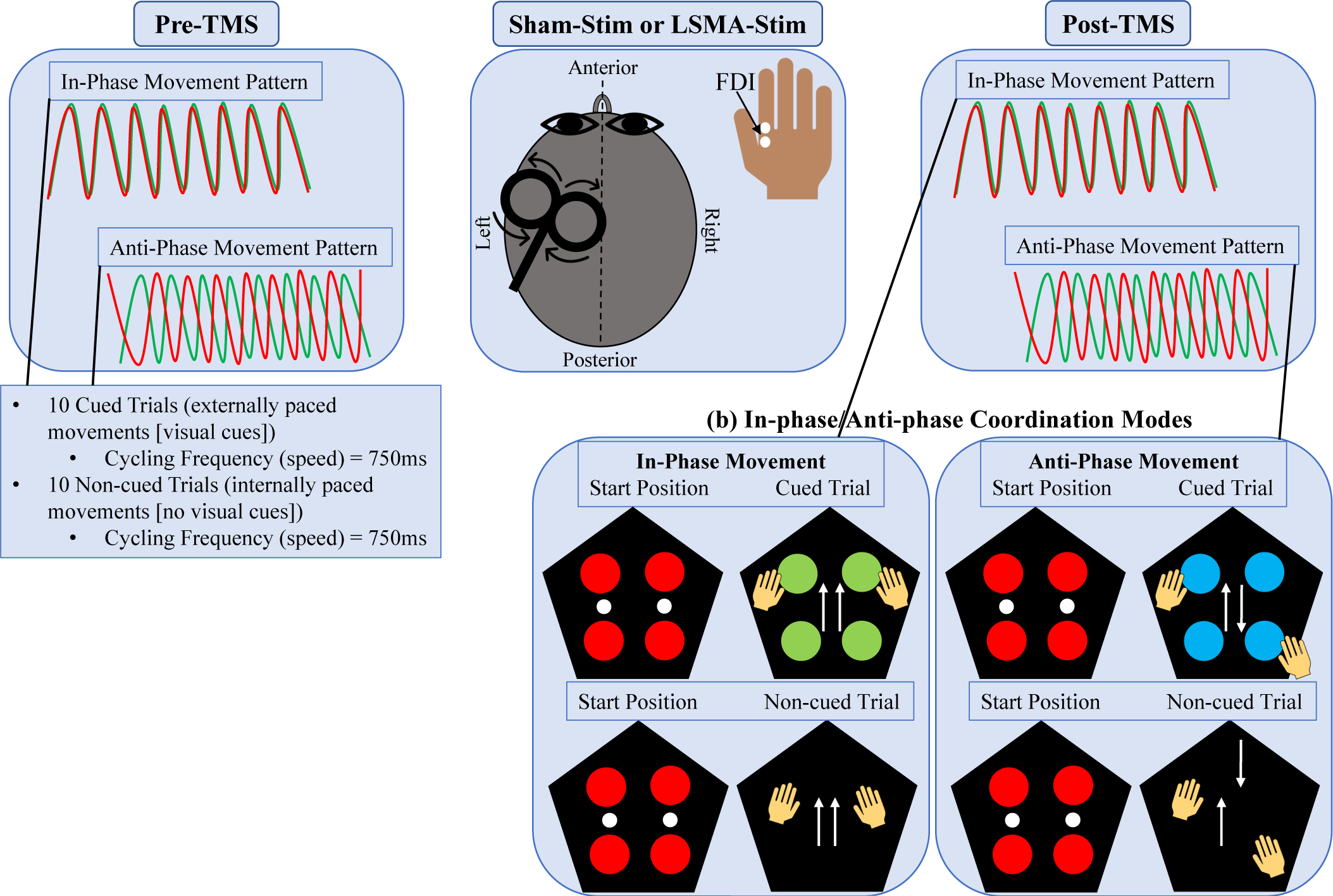
Experimental Setup. (a) Schematic overview of the Pre/Post TMS procedure. The middle panel illustrates two possible stimulation conditions: Sham-Stim and lSMA-Stim. These conditions were randomized across participants. The Pre-TMS and Post-TMS portions of the experiment both consisted of 10 cued trials and 10 non-cued trials, which were randomized. The movement waveforms for in-phase and anti-phase coordination are also displayed. (b) Schematic representation of the two coordination modes: in-phase & anti-phase. Participants were given cued and non-cued trials starting from a red target. During cued trials in the in-phase mode, participants saw flashing green targets and moved their upper limbs in the same direction. During non-cued trials, no visual stimuli were presented, but in-phase movements were still expected. Similarly, during cued trials in the anti-phase mode, participants saw flashing blue targets and moved their upper limbs in the opposite direction. During non-cued trials, no visual stimuli were presented, but anti-phase movements were expected.

During the in-phase or anti-phase coordination modes, trials were divided into cued and non-cued trials. Cued trials provided visual cues to guide the participants’ movements (green targets for in-phase movement and blue targets for anti-phase movement) and were considered externally paced movements, while non-cued trials did not provide visual cues and were considered internally paced movements. A total of 21 trials were completed for both in-phase and anti-phase coordination modes in the pre-TMS session, as well as in the post-TMS session. In each session, a training trial, which served as a cue for participants and acquainted them with the task and movement pace, was conducted initially. However, this trial was not considered in the final analysis, resulting in 20 trials for each coordination mode in both the pre-TMS and post-TMS sessions.

Participants were not informed about the specifics of the cued vs. non-cued trials and were encouraged to perform both types of movements to the best of their ability, particularly during non-cued trials when they did not have any visual cues to guide their movements. Pauses occurred between each trial, and after a 500ms delay, white targets reappeared, signaling the start of the next trial. Approximately 30 oscillations of in-phase and anti-phase movements were performed during each trial.

### Transcranial Magnetic Stimulation Procedure

In this study, we employed a continuous theta burst stimulation (cTBS) paradigm—a form of repetitive transcranial magnetic stimulation (rTMS)—to downregulate targeted regions of the cortex for approximately 20-40 minutes after stimulation, using the Magstim Rapid2 system. The cTBS was delivered in bursts of three pulses at 50Hz, repeated at 200ms intervals, for a total of 600 pulses over 40 seconds at 80% of each participant’s active motor threshold (AMT) (Huang et al., 2005). However, if a participant’s AMT intensity exceeded the safe limit of our equipment, cTBS was administered at the maximum safe intensity of 45% of maximum stimulator output.

To determine the AMT, we administered single pulse TMS to the left primary motor cortex hotspot and recorded visible twitches in the flexed first dorsal interosseous (FDI) muscle in 5 out of 10 trials. Visible twitches were verified by measuring motor-evoked potentials (MEPs) of at least 50 microvolts from the right FDI muscle (Figure 1a). The best location for the motor hotspot was determined by comparing MEP size and consistency at rest. Surface electrode myography (EMG) with Ag/AgCl sintered electrodes over the belly of the right FDI muscle and a ground electrode placed over the bone near the right elbow were used to measure MEPs. Single pulse TMS to the primary motor cortex was delivered using a figure-of-eight coil (Magstim, D702 70mm coil, Carmarthenshire, United Kingdom) held tangential to the scalp surface at an angle of 45^0^ from the anterior-posterior midline, as per standard protocols. See Figure 1a, Sham-Stim or lSMA-Stim panel.

The Magstim Visor2 3-D motion capture-guided neuronavigation system was used to navigate to the lSMA. To achieve this, each participant’s brain model was scaled to the Talairach brain, taking into account head size and shape. Prior literature was consulted to determine the coordinates for lSMA stimulation sites, with the target coordinates set to Talairach −6, −12, 54, as reported by Fabbri et al., 2012. During cTBS, the coil was oriented at 45^0^ from the anterior-posterior midline, facing anterior and held tangential to the scalp, in accordance with the method described by (Janssen et al., 2015). Sham cTBS was administered over the left M1, with the coil facing away from the participant’s head. The experimental sessions were conducted in accordance with the UC Merced IRB protocol for TMS experiments, with a minimum of one day separating each session.

### Data Processing & Analysis

The raw data files containing the hand position data, velocity, and acceleration of each limb of the hand, elbow, and shoulder joints were imported into MATLAB (R2020b, The MathWorks, USA) for offline data processing using the Kinarm MATLAB scripts and custom MATLAB scripts. We calculated both the mean continuous relative phase (ϕ) and the standard deviation of the continuous relative phase (SDϕ). To focus specifically on steady-state performance within each single trial and to compare the cued vs. non-cued trials, we analyzed the data from 12 to 42 seconds per trial for all study participants in both the in-phase and anti-phase coordination modes. It should be noted that during cued and non-cued trials, a different number of oscillations were analyzed for the same 30-second time period as during non-cued trials (no visual targets present), and the pacing of movements was different than when cued trials (visual targets present) were presented.

### Calculation of Continuous Relative Phase (ϕ) and Its Variability (SDϕ)

Continuous relative phase (ϕ) was calculated to quantify and characterize the in-phase and anti-phase coordination modes, along with their variability (Kelso, 1995), using the Kinarm position data from both the left and right hands. The phase angles for each hand were determined using the Hilbert transform approach described by Lamb and Stöckl (Lamb & Stöckl, 2014), which involved amplitude-centering the kinematic signal around zero (Eq. 1) and calculating phase angles using the position at time t, x(t), and their Hilbert transform *H(t) = H(x(t))*.

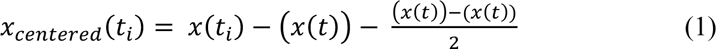

The Hilbert transformation produces a complex analytical signal, ζ(t), with the H(t) of x(t) serving as the imaginary components of the signal. This can be mathematically defined by the following equation (Eq. 2):

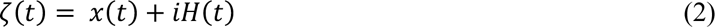

Using the calculation of the complex signal, the phase angle at a given time t_i_ can be determined by calculating the inverse tangent, as shown in the following equation (Eq. 3):

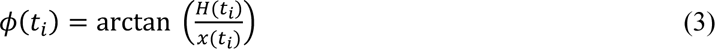

To calculate the ϕ between the right hand (x1(t)) and the left hand (x2(t)), the phase angles of each hand were subtracted from one another. Specifically, the ϕ at time t_i_ was determined using the following equation (Eq. 4), where H1(t) and H2(t) represent the Hilbert transformed signals of the right and left hands, respectively:

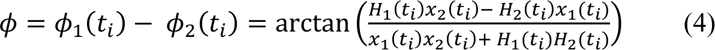

These procedures were applied to all study participants for both the in-phase and anti-phase coordination modes and repeated for each trial. The obtained ϕ values ranged from 0° to 180°, with 0° representing a fully in-phase movement pattern and 180^0^ representing a fully anti-phase movement pattern(Balasubramaniam & Turvey, 2004). In summary, ϕ values of both right and left hand were utilized to quantify and distinguish the two coordination modes, while the standard deviation (SDϕ) of ϕ indicated variability within ϕ.

### Statistical Analyses

Statistical analyses were performed using R (Version 1.3.1093). Linear mixed-effects (LME) regression models were fitted using the lme4 package (Bates et al., 2015). We used LME models to analyze the ϕ and SDϕ, which accounted for the variation in our data contributed by trial number and participant (Gałecki & Burzykowski, 2013). To correct for multiple comparisons, we used Tukey’s method. We extracted estimated marginal means and computed pairwise comparisons with corresponding confidence intervals for cued/non-cued trials, as well as pre- and post-TMS conditions for both in-phase and anti-phase coordination modes using the emmeans R package (Lenth, 2017/2022). Tabular results are presented in Supplementary Table 8, while plots illustrating the pairwise comparisons can be found in Supplementary Figure 2 (see Supplementary materials).

We also fitted multiple LME models to examine potential learning effects from one trial to the next in both coordination modes. We specifically looked at the changes in variability of relative phase, as a decrease is variability is indicative of learning effects (Newell & Vaillancourt, 2001; Schöllhorn et al., 2006; Wu et al., 2014). The results were as follows: During the in-phase coordination mode, no learning effects were observed in the sham stimulation condition (pre-TMS or post-TMS, cued or non-cued). In contrast, there were learning effects observed in the pre-TMS lSMA stimulation (cued trials only), while no learning effects were observed in the non-cued trials. It is worth noting that the observed learning effect was relatively small (β = −0.04), indicating a modest decrease in variability as the trial sequence progressed. In addition, no learning effects were observed for the post-TMS lSMA stimulation (post-TMS). This discrepancy shows that the presence of cues had a selective effect on learning, enhancing overall performance enhancements only while participants obtained specific cues. During the anti-phase coordination mode, no learning effects were observed in the sham (pre-TMS or post-TMS, cued or non-cued) or the lSMA (pre-TMS or post-TMS, cued or non-cued) stimulation condition (refer to Supplementary Figure 1 and Supplementary Tables 6 & 7 for figures and statistical analyses, respectively). The lack of learning effects in this repetitive task may be attributed to the participants’ prior familiarity with the task. Since the participants received training on the task before starting the experimental phase, their previous experience and familiarity with the task may have influenced the observed learning effects. The participants’ prior knowledge and skills acquired during training may have already optimized their performance, leading to a diminished potential for further improvements.

We performed LME model comparisons for both continuous relative phase (ϕ) and variability of continuous relative phase (SDϕ) for in/anti-phase coordination modes to determine the best fit for our data. We found that the main effects model was the most appropriate for ϕ and SDϕ in both coordination modes. The main effects model, which considers the overall influence of predictors on the outcome without accounting for interaction effects, demonstrated superior fit to the data compared to alternative models. This indicates that the overall contribution of predictors was more influential than any potential interactions. By performing model comparisons, we were able to identify important predictors of the outcome variable and assess their relative importance. The significance of predictors was evaluated, allowing us to determine the factors that significantly contributed to the observed coordination modes (Prins & Kingdom, 2018). Supplementary Tables 2-5 present the detailed results of the model comparisons for ϕ and SDϕ in both in-phase and anti-phase coordination modes.

The best fit model was determined using five LME equations for ϕ and SDϕ (refer to Supplementary Table 1 for the equations). Model 1 served as a null model with no predictors. Model 2 used only the main effects of stimulation type (lSMA vs. Sham) and trial type (cued vs. non-cued). Model 3 incorporated the main effects of stimulation type and trial type, along with the two-way interactions of stimulation type and trial type. Model 4 was a model with all possible two-way interactions. Model 5 was a three-way interaction between pre/post session, stimulation type, and trial type. All models included trial number and participants as random effects. LME models with random intercepts such as the ones above take into account the variability in individual subjects. This is important as it allows for generalizability and helps avoid biased estimates of model parameters, as it allows the model to capture the underlying variance and covariance more accurately in the data.

## Results

### Continuous Relative Phase (ϕ)

Figures 2 and 3 illustrate the changes in continuous relative phase between pre- and post-TMS stimulation during the in-phase and anti-phase coordination mode. In Figure 2, the average relative phase values for each subject are shown before and after sham or lSMA stimulation, across cued and non-cued trials. Note that the closer participants are to 0^0^ during in-phase coordination mode and 180^0^ during anti-phase coordination mode, the more stable their movement.

**Figure 2.**
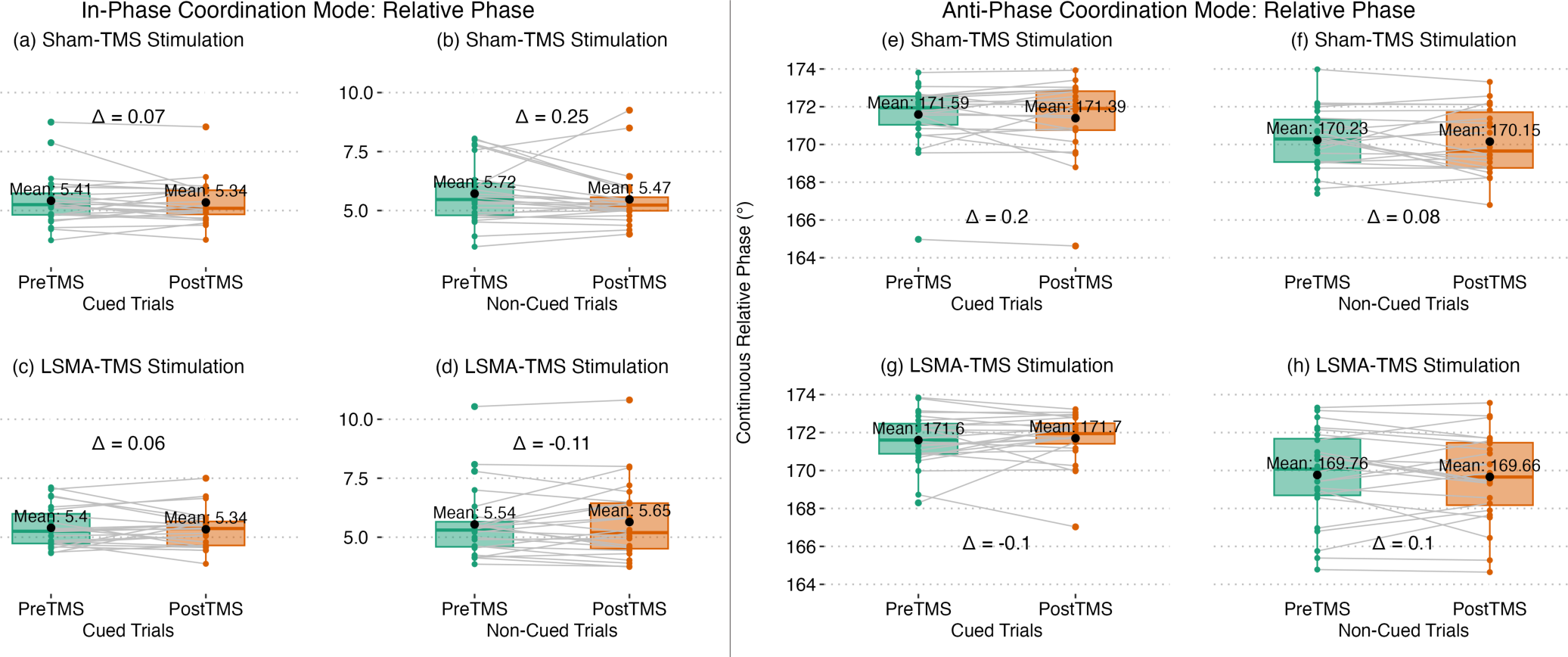
Pre- and post-TMS changes in continuous relative phase (ϕ) during sham and lSMA stimulation for cued and non-cued trials, averaged across trials for each subject. The black dot in each figure represents the mean. Panels (a-d) depict the relative phase changes during the in-phase coordination mode, while panels (e-h) show the changes during anti-phase coordination mode. Specifically, the closer the relative phase values are to 0^0^ for in-phase movements and 180^0^ for anti-phase movements, the more stable the movements are.

**Figure 3.**
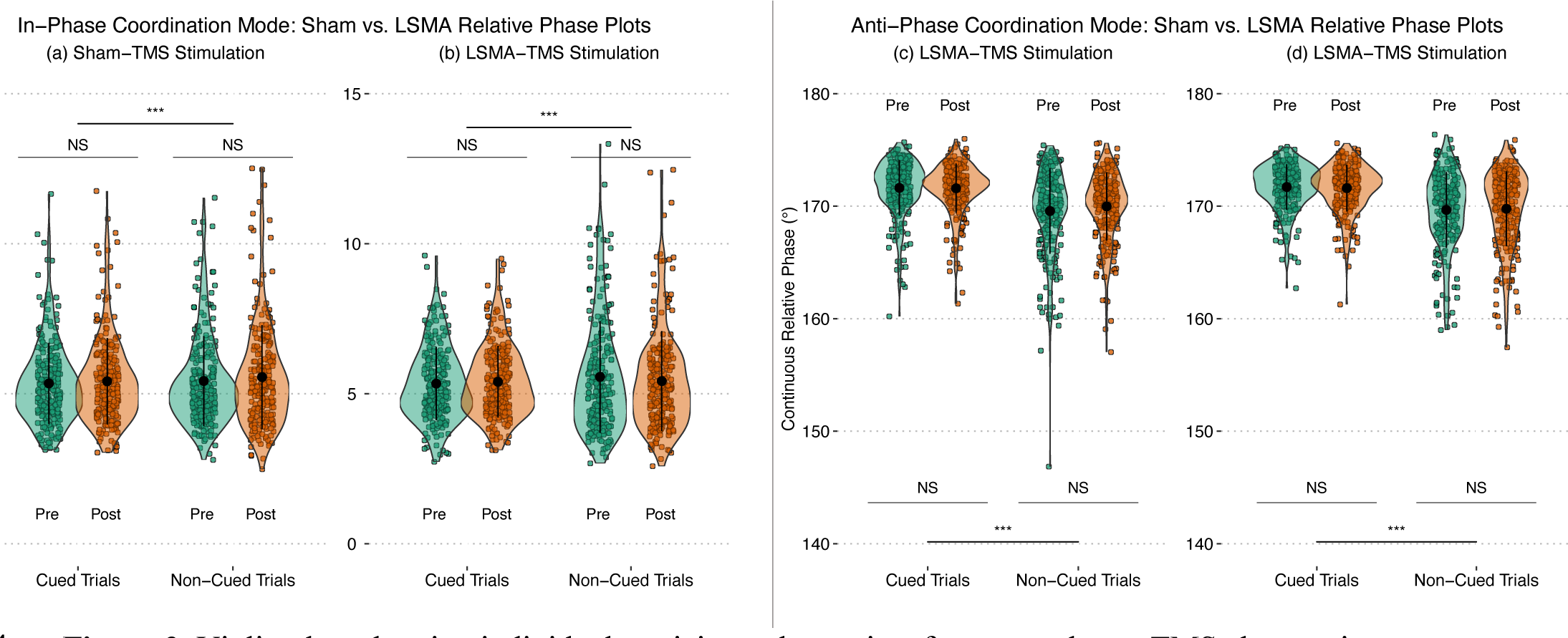
Violin plots showing individual participant data points for pre- and post-TMS changes in continuous relative phase (ϕ) during sham and lSMA stimulation for cued and non-cued trials. The width of each violin represents the density of data points, and the black dot inside indicates the mean. Panels (a-b) display data for the in-phase coordination mode, while panels (c-d) show data for the anti-phase coordination mode. Note that there is a significant difference between cued and non-cued trials for both coordination modes, but no significant differences were found between pre- and post-TMS for either sham or lSMA stimulation.

As can be seen during the in-phase coordination mode, for Figures 2a & b (sham-TMS), there is an average change in relative phase between pre- and post-TMS of 0.07^0^ for cued trials and 0.25^0^ for non-cued trials, signifying a reduction in relative phase following sham-TMS. Conversely, in figures 2c & d (lSMA-TMS) there is an average change in relative phase between pre- and post-TMS stimulation of 0.06^0^ for cued trials and −0.11^0^ for non-cued trials. These results indicate a decrease in relative phase during cued trials and an increase or worsening of relative phase during non-cued trials.

In anti-phase coordination mode, Figure 2e and 2f (sham-TMS) demonstrate an average change in relative phase between pre- and post-TMS of 0.2^0^ and 0.08^0^ for cued and non-cued trials, respectively, signifying a worsening of relative phase. Meanwhile, Figure 2g and h (lSMA-TMS) show an average change in relative phase between pre- and post-TMS of −0.1^0^ and 0.1^0^ for cued and non-cued trials, respectively, indicating slight improvement during cued trials and slight worsening of movement after lSMA stimulation.

To quantify the differences between movements patterns across different stimulation conditions in both the in-phase and anti-phase coordination modes, LME model were implemented, with fixed effects of pre/post-TMS, stimulation conditions (sham/lSMA), and trial type (cued/non-cued) and random effects of participants and trials to account for variance in the data. Based on model comparisons, both the in-phase and anti-phase coordination models included the main effect only (see Model 2 in Supplementary Table 1). The results for continuous relative phase and variability of continuous relative phase can be found in Table 1.

**Table 1.** Linear mixed-effects model (LME) analysis results for continuous relative phase (ϕ) during in-phase and anti-phase coordination modes. The table presents the effects of pre- and post-TMS stimulation during both sham and lSMA conditions, as well as cued and non-cued trials. Note: Statistical significance of p<0.05 after Tukey’s multiple comparisons test is indicated in bold.

| Mean Continuous Relative Phase ($\phi$) | | | | | | | |
| --- | --- | --- | --- | --- | --- | --- | --- |
| In-phase Coordination Mode |  |  |  | Anti-phase Coordination Mode |  |  |  |
| Fixed effects | $\beta$ | SE | $p(\chi^2)$ | Fixed effects | $\beta$ | SE | $p(\chi^2)$ |
| Intercept | 5.33 | 0.18 | **<0.001** | Intercept | 171.58 | 0.35 | **<0.001** |
| Pre/Post [PreTMS] | 0.01 | 0.07 | 0.325 | Pre/Post [PreTMS] | 0.09 | 0.11 | 0.391 |
| Stimulation type [Sham] | 0.01 | 0.07 | 0.934 | Stimulation type [Sham] | 0.00 | 0.11 | 0.964 |
| Trial type [non-cued] | 0.22 | 0.07 | **0.001** | Trial type [non-cued] | -1.89 | 0.11 | **<0.001** |
| Random effects | Groups |  | SD | Random effects | Groups |  | SD |
|  | Participant | Intercept | 0.80 |  | Participant | Intercept | 1.61 |
|  | Trial | Intercept | 0.08 |  | Trial | Intercept | 0.00 |
|  | Residual |  | 1.53 |  | Residual |  | 2.37 |
|  | Observations: 1920 |  |  |  | Observations: 1920 |  |  |

### Variability of Continuous Relative Phase (SDϕ)

To access variability (SDϕ), we utilized a linear mixed effects model (specifically, Model 2 in Supplementary Table 1) to analyze both in-phase and anti-phase coordination modes. This model incorporated fixed defects of pre/post-TMS, stimulation conditions (sham/lSMA), and trial type (cued/non-cued), while accounting for random effects of participants and trials to address variance in the data. Our findings demonstrated statistically significant differences between cued and non-cued trial types in both the in-phase coordination mode (Figure 5a and 5b) and anti-phase coordination mode (Figure 5c and 5d). However, no significant differences were observed in pre- or post-stimulation or stimulation conditions (sham/lSMA).

**Figure 4.**
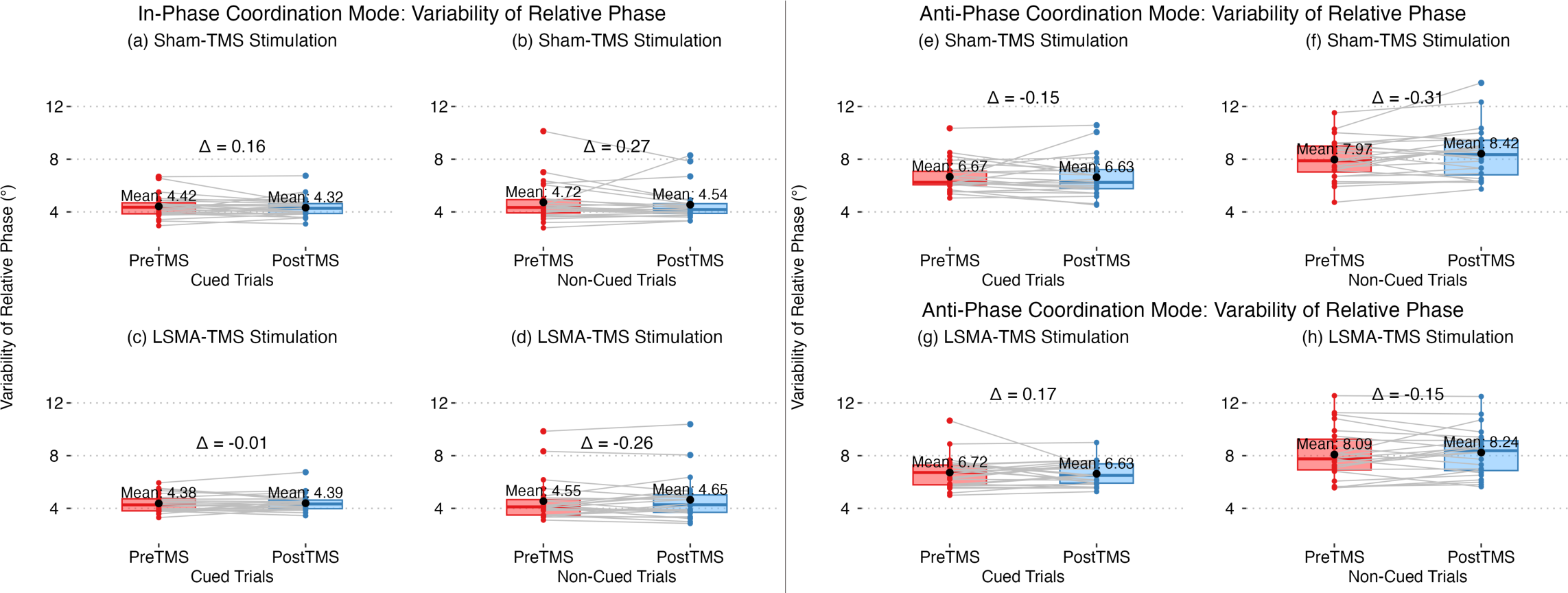
Pre- and post-TMS changes in variability of continuous relative phase (SDϕ) during sham and lSMA stimulation for cued and non-cued trials, averaged across trials for each subject. The black dot signifies the mean. Panels (a-d) indicate the variability changes during the in-phase coordination mode, whereas panels (e-h) show the changes during anti-phase coordination mode.

**Figure 5.**
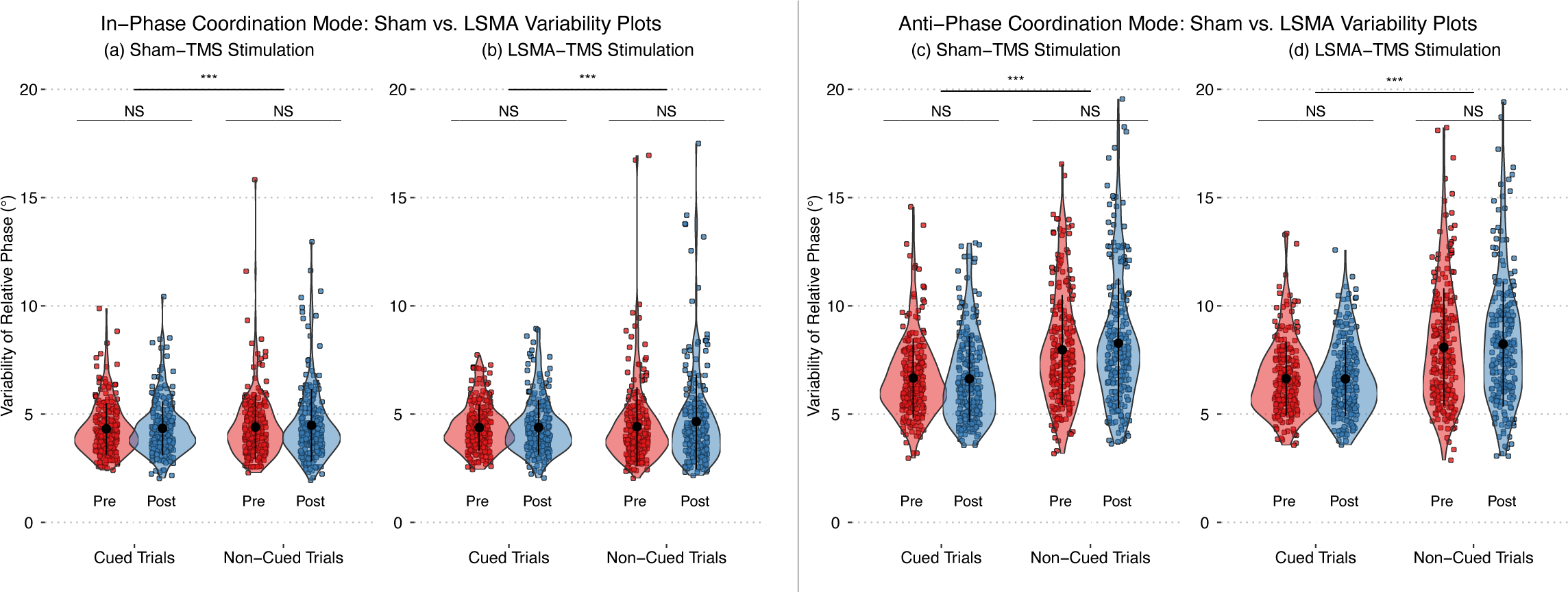
Violin plots illustrating individual participant data points for pre- and post-TMS changes in variability of continuous relative phase SDϕ) during sham and lSMA stimulation for cued and non-cued trials. Each violin plot represents individual participate data points, with the width indicating the data density, and the black dot indicating the mean. Panels (a-b) display data for the in-phase coordination mode, while panels (c-d) show data for the anti-phase coordination mode. Note that there is a significant difference between cued and non-cued trials for both coordination modes, but no significant differences were found between pre- and post-TMS for either sham or lSMA stimulation.

Furthermore, although no statically significant differences were observed between the pre- and post-stimulation conditions, Figure 4 highlights subtle variations in the average relative phase between two conditions for variability in both in-phase and anti-phase coordination modes. Specifically, in the in-phase coordination mode, Figure 4a (sham stimulation) shows an average variability change of 0.16^0^ between pre- and post-stimulation in non-cued trials, while Figure 4b (sham stimulation), displays an average variability change of 0.27^0^ in cued trials. Similarly, Figure 4c (lSMA stimulation) depicts an average variability change of −0.01^0^ between pre- and post-stimulation in cued trials, whereas Figure 4d (lSMA stimulation) indicates the average variability change of −0.26^0^, indicating a slight increase in variability following lSMA stimulation.

During the anti-phase coordination model, Figure 4e (sham stimulation) presents an average variability change of −0.15^0^ between pre- and post-stimulation in cued trials, whereas Figure 4f (sham stimulation) indicates an average variability change of −0.31^0^ in non-cued trials. Additionally, Figure 4g (lSMA stimulation) displays an average variability change of 0.17^0^ between the pre- and post-stimulation in cued trials, and Figure 4h (lSMA stimulation) indicates an average variability change of −0.15^0^ in non-cued trials, again suggesting a slight increase in variability following lSMA stimulation. Table 2 depict the variability results of the linear mixed effects model for in-phase and anti-phase movement patterns, respectively.

**Table 2.** Results of Linear Mixed-Effects Model (LME) Analysis for Variability of Continuous Relative Phase (SDϕ) in In-Phase and Anti-Phase Coordination Modes. The table presents the effects of pre- and post-TMS stimulation under sham and lSMA conditions, as well as cued and non-cued trials. Note: Statistical significance at p<0.05 after Tukey’s multiple comparisons test is indicated in bold font.

| Variability of Continuous Relative Phase ($SD\phi$) | | | | | | | |
| --- | --- | --- | --- | --- | --- | --- | --- |
| In-phase Coordination Mode |  |  |  | Anti-phase Coordination Mode |  |  |  |
| Fixed effects | $\beta$ | SE | $p(\chi^2)$ | Fixed effects | $\beta$ | SE | $p(\chi^2)$ |
| Intercept | 4.35 | 0.16 | <0.001 | Intercept | 6.72 | 0.25 | <0.001 |
| Pre/Post [PreTMS] | 0.04 | 0.09 | 0.655 | Pre/Post [PreTMS] | -0.12 | 0.10 | 0.249 |
| Stimulation type [Sham] | 0.01 | 0.09 | 0.936 | Stimulation type [Sham] | 0.00 | 0.10 | 0.988 |
| Trial type [non-cued] | 0.24 | 0.09 | 0.006 | Trial type [non-cued] | 1.52 | 0.10 | <0.001 |
| Random effects | Groups |  | SD | Random effects | Groups |  | SD |
|  | Participant | Intercept | 0.68 |  | Participant | Intercept | 1.10 |
|  | Trial | Intercept | 0.05 |  | Trial | Intercept | 0.00 |
|  | Residual |  | 1.91 |  | Residual |  | 2.21 |
|  | Observations: 1920 |  |  |  | Observations: 1920 |  |  |

## Discussion

The current experiment investigated the impact of continuous theta burst stimulation (cTBS) targeting the supplementary motor area (SMA) on in-phase and anti-phase coordination modes in a bimanual coordination task. This study builds upon previous research on bimanual coordination tasks, focusing on the influence of SMA stimulation during in/anti-phase coordination (Shima & Tanji, 1998; Thickbroom et al., 2000; Obhi et al., 2002; Serrien et al., 2002; Chen et al., 2005; Steyvers et al., 2003; Neva et al., 2014). The study employed a steady-state system of coordination dynamics, exclusively examining true in/anti-phase movements and disregarding phase transitions. The mean continuous relative phase (ϕ) and variability in relative phase (SDϕ) were assessed during both the in/anti-phase coordination modes between cued and non-cued trials in both pre- and post-TMS stimulation conditions (sham and lSMA). A total of 11 cued and 10 non-cued trials were completed for pre-TMS stimulation (for both sham & lSMA stimulation conditions). Similarly, 11 cued and 10 non-cued trials were completed for both in/anti-phase coordination modes for post-TMS stimulation (for both sham & lSMA stimulation conditions). The first cued trial was excluded from the analysis, resulting in a total of 20 trials for both pre-TMS (sham and left SMA) and post-TMS (sham and left SMA) conditions.

The results revealed significant differences between cued and non-cued conditions for both ϕ: in-phase ϕ (β = 0.22; *SE* = 0.07; *p* = 0.001); anti-phase ϕ (β = −1.89; *SE* = 0.11; *p* < 0.001) and SDϕ: in-phase SDϕ (β = 0.24; *SE* = 0.09; *p* = 0.006); anti-phase SDϕ (β = 1.52; *SE* = 0.10; *p* < 0.001). This indicates that visual cues did have a significant influence on participants’ ability to perform accurate and synchronized bimanual movements. These findings regarding the effect of visual cues are consistent with previous research indicating that external cues, such as visual stimuli, may help enhance bimanual coordination (Debaere et al., 2003). Visual cues provide temporal and spatial information that assists participants in synchronizing their movements and maintaining a stable bimanual pattern. In the present study, the presence of visual cues likely facilitated participants’ utilization of their internal sense of timing and spatial coordination, resulting in improved performance in the cued condition.

Contrary to our expectations, the downregulation of the lSMA through cTBS did not lead to significant impairments in bimanual coordination performance. Thus, these findings suggest that lSMA stimulation does not have a significant effect during bimanual in/anti-phase movement, at least in the context of the present experiment. It’s important to note that this finding does not discount the potential involvement of the SMA in bimanual coordination, as the lack of significant effects could be influenced by various factors. Thus, this prompts the exploration of potential reasons for the lack of observed effects. Several factors could contribute to the absence of significant effects, including methodological limitations, individual differences, and functional redundancy within the motor system (Wassermann & Lisanby, 2001; Balasubramaniam, 2013; Gröhn et al., 2019; Turi et al., 2021; Hanlon & McCalley, 2022). Exploring these limitations is important in order to gain a comprehensive understanding of the findings.

First, the lack of significant results may be partly due to methodological limitations. Factors such as the choice of stimulation parameters, including cTBS intensity, duration, and frequency, may influence the efficacy of SMA modulation (Bestmann & Krakauer, 2015; Huang et al., 2005). Variability in stimulation parameters across studies may lead to inconsistent outcomes and hinder the detection of significant effects. Additionally, the timing of stimulation relative to task execution and the duration of the stimulation effects may influence the outcome. The SMA is involved in motor planning and execution, thus the specific timing of cTBS relative to these processes may influence its effects. For instance, cTBS applied before a motor task may have different consequences compared to stimulation administer during the task (Toyokura et al., 2002; Steyvers et al., 2003). Moreover, the duration of the stimulation effects plays a significant role in understanding the temporal dynamics of SMA modulation. Thus, variations in these factors across studies could contribute to inconsistencies in results.

Second, individual differences among participants may have influenced the response to SMA stimulation. Inter-individual variability in neuroanatomy, functional connectivity, and baseline motor performance could impact the effectiveness of cTBS on bimanual coordination (Reis et al., 2009; Lage-Castellanos et al., 2010). The SMA is a heterogeneous region with distinct subregions that serve different functions (Nachev et al., 2008). Variability in the exact stimulation site within the SMA might lead to differences in responses across individuals. Moreover, variations in the recruitment of compensatory mechanisms or alternative neural pathways for bimanual coordination may further contribute to the individual differences observed (Ward & Frackowiak, 2006). The brain exhibits remarkable plasticity, allowing it to adapt and undergo functional changes in response to injury or functional demands (Péran et al., 2014). Thus, some individuals may rely more on compensatory mechanisms or engage alternative neural pathways enabling the ability to move in/anti-phase bimanually. These individual differences in neural plasticity and compensatory strategies could influence the response to SMA stimulation.

Third, the lack of significant effects could be attributed to functional redundancy within the motor system. This redundancy allows for the motor system to adapt to changes in the environment, compensate for injury, and perform skilled movements efficiently. The SMA is one component of a larger network involved in motor control, including the primary motor cortex, premotor cortex, and cerebellum (Picard & Strick, 1996; Bracewell et al., 2005). These regions exhibit overlapping functionality, with the potential for compensation in the absence of SMA modulation. Further evidence suggests that neural plasticity and reorganization can also occur at the network level rather than being solely dependent on SMA functionality. For example, a review by Bestmann & Krakauer discussed the concept of distributed motor networks and their ability to adapt in response to perturbations. They emphasized that motor functions are not solely localized to specific brain regions, but rather involves a more distributed network that can dynamically reorganize in order to maintain optical motor performance (Bestmann & Krakauer, 2015). Considering the redundancy and flexibility in motor networks, it seems plausible that cTBS of the SMA region along may not yield significant effects of certain motor tasks and other motor regions may compensate for the perturbation, maintaining adequate motor performance.

It is also important to consider the limitations of the study itself. The sample size and statistical power may have influenced the ability to detect significant effects (Héroux et al., 2015; Mitra et al., 2019). The study design and the specific tasks employed for assessing bimanual coordination could have influenced the sensitivity to detect changes induced by SMA stimulation. Additionally, the chosen outcome measures may not have been sensitive enough to capture subtle alterations in bimanual coordination resulting from SMA modulation. To gain a deeper understanding of the factors contributing to the lack of observed effects, future research should consider optimizing the stimulation parameters, inclusion of larger and more diverse sample sizes, and the utilization of more refined and sensitive outcome measures.

Overall, the present experiment underscores the importance of visual cues in facilitating bimanual coordination. It provides evidence that the presence or absence of visual cues significantly impacts a participants’ ability to perform accurate and synchronized bimanual movements. However, it did not find a significant role for the lSMA in bimanual coordination, as the downregulation of this region did not lead to observable impairments by way of ϕ and SDϕ. These results suggest that the lSMA may not be a critical component specifically involved in the coordination of bimanual movements or that other brain regions may compensate for its temporary inhibition. Further investigation is needed in order to understand the precise contributions of the SMA in bimanual coordination task. Future studies could explore potential methodological improvements to enhance the sensitivity of stimulation protocols, such as considering alternative parameters or different stimulation techniques. Moreover, it is crucial to examine the interactions between the SMA and other brain regions involved in motor control. Functional redundancy within the motor system may have masked the effects of lSMA downregulation in this study. Investigating the interplay between the SMA and regions such as the primary motor cortex, premotor cortex, and cerebellum could provide valuable insights into the compensatory mechanisms and distributed neural networks underlying bimanual coordination.

## Notes

### Competing Interest Statement

The authors have declared no competing interest.

