## Supplementary Materials for "cTBS Over the Left Supplementary Motor Area (lSMA) Does Not Modulate Rhythmic Bimanual Coordination in the Presence and Absence of Visual Cues"

**Supplementary Table 1**. Linear Mixed Effects Model Equations for Model Comparisons in In/Anti-Phase Coordination Modes.

| Model Type | Equation |
|---|---|
| Model 1:<br>Null model with no predictors: | $\phi$ or SD$\phi$ ~ N ($\alpha_{j[i],k[i]}$, $\sigma^2$)<br>$\alpha_j$ ~ N ($\mu_{\alpha j}$, $\sigma^2_{\alpha j}$), for Participant j = 1,…,J<br>$\alpha_k$ ~ N ($\mu_{\alpha k}$, $\sigma^2_{\alpha k}$), for Trial number k = 1,…,K |
| Model 2:<br>Main effects model | $\phi$ or SD$\phi$ ~ N ($\mu$, $\sigma^2$)<br>$\mu = \alpha_{j[i],k[i]} + \beta_1$(Pre or Post) + $\beta_2$(Stim. Type) + $\beta_3$(Trial type)<br>$\alpha_j$ ~ N ($\mu_{\alpha j}$, $\sigma^2_{\alpha j}$), for Participant j = 1,…,J<br>$\alpha_k$ ~ N ($\mu_{\alpha k}$, $\sigma^2_{\alpha k}$), for Trial number k = 1,…,K |
| Model 3:<br>Main effects and two-way interaction model | $\phi$ or SD$\phi$ ~ N ($\mu$, $\sigma^2$)<br>$\mu = \alpha_{j[i],k[i]} + \beta_1$(Pre or Post) + $\beta_2$(Stim. Type) + $\beta_3$(Trial type)<br>+ $\beta_4$(Pre or Post x Stimulation type)<br>$\alpha_j$ ~ N ($\mu_{\alpha j}$, $\sigma^2_{\alpha j}$), for Participant j = 1,…,J<br>$\alpha_k$ ~ N ($\mu_{\alpha k}$, $\sigma^2_{\alpha k}$), for Trial number k = 1,…,K |
| Model 4:<br>All possible two-way interactions | $\phi$ or SD$\phi$ ~ N ($\mu$, $\sigma^2$)<br>$\mu = \alpha_{j[i],k[i]} + \beta_1$(Pre or Post) + $\beta_2$(Stim. Type) + $\beta_3$(Trial type)<br>+ $\beta_4$(Pre or Post x Stim. Type) + $\beta_5$(Pre or Post x Trial type)<br>+ $\beta_6$(Stim. Type x Trial type)<br>$\alpha_j$ ~ N ($\mu_{\alpha j}$, $\sigma^2_{\alpha j}$), for Participant j = 1,…,J<br>$\alpha_k$ ~ N ($\mu_{\alpha k}$, $\sigma^2_{\alpha k}$), for Trial number k = 1,…,K |
| Model 5:<br>Full model with three-way interaction | $\phi$ or SD$\phi$ ~ N ($\mu$, $\sigma^2$)<br>$\mu = \alpha_{j[i],k[i]} + \beta_1$(Pre or Post) + $\beta_2$(Stim. Type) + $\beta_3$(Trial type)<br>+ $\beta_4$(Pre or Post x Stim. Type) + $\beta_5$(Pre or Post x Trial type)<br>+ $\beta_6$(Stim. Type x Trial type) + $\beta_7$(Pre or Post x Stim. Type x Trial type)<br>$\alpha_j$ ~ N ($\mu_{\alpha j}$, $\sigma^2_{\alpha j}$), for Participant j = 1,…,J<br>$\alpha_k$ ~ N ($\mu_{\alpha k}$, $\sigma^2_{\alpha k}$), for Trial number k = 1,…,K |

**Supplementary Table 2.** Linear mixed effects model comparisons for continuous relative phase (ϕ) during in-phase coordination mode.

| **Model**<br>**Continuous Relative Phase (ϕ) –**<br>**In-phase Coordination Mode** | **df** | **AIC** | **BIC** | **χ2 likelihood ratio** | **χ2 p-value** | **Model Comparison** |
|---|---|---|---|---|---|---|
| Model 1: Null model with no predictors | | 7166.9 | 7189.1 | | | |
| Model 2: Main effects only model | 3 | 7161.7 | 7200.6 | 11.25 | **0.010*** | Model 1 vs. Model 2 |
| Model 3: Main effect and two-way interaction model | 1 | 7162.0 | 7206.5 | 1.61 | 0.204 | Model 2 vs. Model 3 |
| Model 4: All possible two-way interaction model | 2 | 7166.0 | 7221.6 | 0.01 | 0.993 | Model 3 vs. Model 4 |
| Model 5: Three-way interaction model | 1 | 7166.6 | 7227.8 | 1.39 | 0.239 | Model 4 vs. Model 5 |
| *Final model equation: Model 2 (Main effects only model)* | | | | | | |

**Supplementary Table 3.** Linear mixed effects model comparisons for variability of relative phase (SDϕ) during in-phase coordination mode.

| **Model**<br>**Variability of Continuous Relative Phase (ϕ) –**<br>**In-phase Coordination Mode** | **df** | **AIC** | **BIC** | **χ2 likelihood ratio** | **χ2 p-value** | **Model Comparison** |
|---|---|---|---|---|---|---|
| Model 1: Null model with no predictors | | 7996.1 | 8081.3 | | | |
| Model 2: Main effects only model | 3 | 7994.2 | 8033.2 | 7.84 | **0.049*** | Model 1 vs. Model 2 |
| Model 3: Main effect and two-way interaction model | 1 | 7995.0 | 8039.4 | 1.29 | 0.256 | Model 2 vs. Model 3 |
| Model 4: All possible two-way interaction model | 2 | 7998.9 | 8054.5 | 0.09 | 0.955 | Model 3 vs. Model 4 |
| Model 5: Three-way interaction model | 1 | 8000.6 | 8061.8 | 0.25 | 0.620 | Model 4 vs. Model 5 |
| *Final model equation: Model 2 (Main effects only model)* | | | | | | |

**Supplementary Table 4.** Linear mixed effects model comparisons for continuous relative phase (ϕ) during anti-phase coordination mode.

| **Model**<br>**Continuous Relative Phase (ϕ) –**<br>**Anti-phase Coordination Mode** | **df** | **AIC** | **BIC** | **χ2 likelihood ratio** | **χ2 p-value** | **Model Comparison** |
|---|---|---|---|---|---|---|
| Model 1: Null model with no predictors | | 9142.2 | 9164.4 | | | |
| Model 2: Main effects only model | 3 | 8865.9 | 8904.8 | 282.25 | **<2e-16*** ** | Model 1 vs. Model 2 |
| Model 3: Main effect and two-way interaction model | 1 | 8867.2 | 8911.6 | 0.74 | 0.390 | Model 2 vs. Model 3 |
| Model 4: All possible two-way interaction model | 2 | 8868.7 | 8924.3 | 2.46 | 0.292 | Model 3 vs. Model 4 |
| Model 5: Three-way interaction model | 1 | 8870.4 | 8931.6 | 0.32 | 0.574 | Model 4 vs. Model 5 |
| *Final model equation: Model 2 (Main effects only model)* | | | | | | |

**Supplementary Table 5.** Linear mixed effects model comparisons for variability of relative phase (SDϕ) during anti-phase coordination mode.

| **Model**<br>**Variability of Continuous Relative Phase (ϕ) –**<br>**Anti-phase Coordination Mode** | **df** | **AIC** | **BIC** | **χ2 likelihood ratio** | **χ2 p-value** | **Model Comparison** |
|---|---|---|---|---|---|---|
| Model 1: Null model with no predictors | | 8788.5 | 8810.7 | | | |

| | | | | | | |
|---|---|---|---|---|---|---|
| Model 2: Main effects only model | 3 | 8579.9 | 8618.8 | 214.64 | **<2e-16*** ** | Model 1 vs. Model 2 |
| Model 3: Main effect and two-way interaction model | 1 | 8581.1 | 8625.6 | 0.77 | 0.380 | Model 2 vs. Model 3 |
| Model 4: All possible two-way interaction model | 2 | 8581.7 | 8637.3 | 3.35 | 0.187 | Model 3 vs. Model 4 |
| Model 5: Three-way interaction model | 1 | 8583.4 | 8644.5 | 0.37 | 0.545 | Model 4 vs. Model 5 |
| *Final model equation: Model 2 (Main effects only model)* | | | | | | |

**Supplementary Figure 1.**

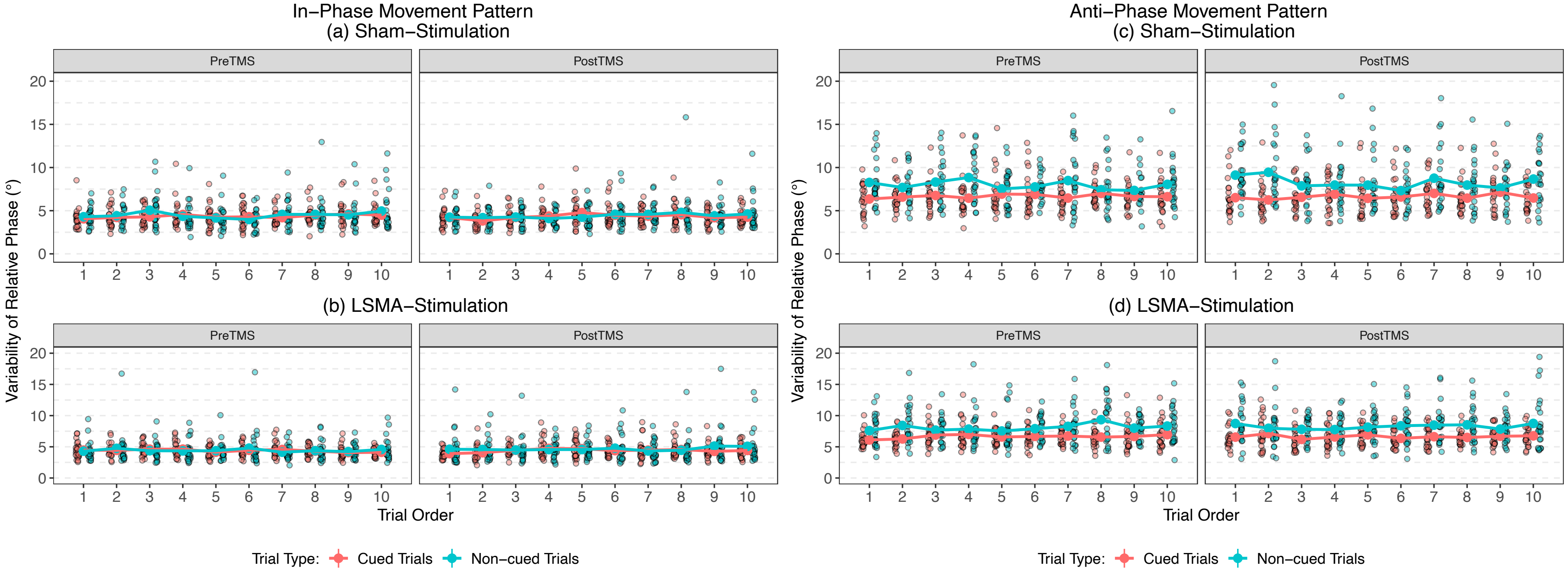

**S1.** Serial plots illustrating the variability of relative phase behavior during pre- and post-TMS under Sham and lSMA stimulation for both in-phase (a-b) and anti-phase (c-d) coordination modes. The plots present trials averaged across all 24 participants.

**Supplementary Table 6.** Linear Mixed Effects Models Investigating Learning Effects in In-Phase Coordination Mode.

| Trial type | Effect of Serial Order & Trial Type on Variability of Relative Phase (SDϕ) In-Phase Coordination Mode | | | | | | | |
|---|---|---|---|---|---|---|---|---|
| | **Sham Stimulation** | | | | | | | |
| | **Pre-TMS** | | | | **Post-TMS** | | | |
| **Cued Trials** | **Fixed Effects** | **β** | **SE** | **p($\chi^2$)** | **Fixed Effects** | **β** | **SE** | **p($\chi^2$)** |
| | Intercept | 4.26 | 0.26 | **<0.001** | Intercept | 4.18 | 0.19 | **<0.001** |
| | Serial Order | 0.03 | 0.03 | 0.406 | Serial Order | 0.03 | 0.02 | 0.237 |
| | **Random effects** | **Groups** | | **SD** | **Random effects** | **Groups** | | **SD** |
| | | Participant | Intercept | 0.74 | | Participant | Intercept | 0.66 |
| | | Residual | | 1.49 | | Residual | | 0.99 |
| | | Observations: 240 | | | | Observations: 240 | | |
| **Non-cued Trials** | **Fixed Effects** | **β** | **SE** | **p($\chi^2$)** | **Fixed Effects** | **β** | **SE** | **p($\chi^2$)** |
| | Intercept | 4.30 | 0.46 | **<0.001** | Intercept | 4.36 | 0.34 | **<0.001** |
| | Serial Order | 0.08 | 0.08 | 0.229 | Serial Order | 0.03 | 0.04 | 0.413 |
| | **Random effects** | **Groups** | | **SD** | **Random effects** | **Groups** | | **SD** |
| | | Participant | Intercept | 1.21 | | Participant | Intercept | 1.11 |
| | | Residual | | 2.82 | | Residual | | 1.85 |
| | | Observations: 240 | | | | Observations: 240 | | |
| | **lSMA Stimulation** | | | | | | | |
| | **Pre-TMS** | | | | **Post-TMS** | | | |
| **Cued Trials** | **Fixed Effects** | **β** | **SE** | **p($\chi^2$)** | **Fixed Effects** | **β** | **SE** | **p($\chi^2$)** |
| | Intercept | 4.62 | 0.17 | **<0.001** | Intercept | 4.21 | 0.20 | **<0.001** |
| | Serial Order | -0.04 | 0.02 | **0.026** | Serial Order | 0.03 | 0.02 | 0.178 |
| | **Random effects** | **Groups** | | **SD** | **Random effects** | **Groups** | | **SD** |
| | | Participant | Intercept | 0.62 | | Participant | Intercept | 0.58 |
| | | Residual | | 0.85 | | Residual | | 1.11 |
| | | Observations: 240 | | | | Observations: 240 | | |
| **Non-cued Trials** | **Fixed Effects** | **β** | **SE** | **p($\chi^2$)** | **Fixed Effects** | **β** | **SE** | **p($\chi^2$)** |
| | Intercept | 4.39 | 0.42 | **<0.001** | Intercept | 4.41 | 0.38 | **<0.001** |
| | Serial Order | 0.03 | 0.05 | 0.565 | Serial Order | 0.04 | 0.03 | 0.180 |
| | **Random effects** | **Groups** | | **SD** | **Random effects** | **Groups** | | **SD** |
| | | Participant | Intercept | 1.39 | | Participant | Intercept | 1.58 |

| | | | | |
|---|---|---|---|---|
| | Residual<br>Observations:<br>240 | 2.21 | Residual<br>Observations:<br>240 | 1.48 |

**Supplementary Table 7.** Linear Mixed Effects Models Investigating Learning Effects in the Anti-Phase Coordination Mode.

| Trial type | Effect of Serial Order & Trial Type on Variability of Relative Phase (SDϕ) Anti-Phase Coordination Mode | | | | | | | |
|---|---|---|---|---|---|---|---|---|
| | **Sham Stimulation** | | | | | | | |
| | **Pre-TMS** | | | | **Post-TMS** | | | |
| **Cued Trials** | **Fixed Effects** | **β** | **SE** | **p($\chi^2$)** | **Fixed Effects** | **β** | **SE** | **p($\chi^2$)** |
| | Intercept | 6.53 | 0.30 | **<0.001** | Intercept | 6.44 | 0.35 | **<0.001** |
| | Serial Order | 0.02 | 0.03 | 0.456 | Serial Order | 0.04 | 0.03 | 0.256 |
| | **Random effects** | **Groups** | | **SD** | **Random effects** | **Groups** | | **SD** |
| | | Participant | Intercept | 1.07 | | Participant | Intercept | 1.42 |
| | | Residual | | 1.47 | | Residual | | 1.37 |
| | | Observations: 240 | | | | Observations: 240 | | |
| **Non-cued Trials** | **Fixed Effects** | **β** | **SE** | **p($\chi^2$)** | **Fixed Effects** | **β** | **SE** | **p($\chi^2$)** |
| | Intercept | 8.29 | 0.41 | **<0.001** | Intercept | 8.53 | 0.54 | **<0.001** |
| | Serial Order | -0.06 | 0.05 | 0.223 | Serial Order | -002 | 0.07 | 0.751 |
| | **Random effects** | **Groups** | | **SD** | **Random effects** | **Groups** | | **SD** |
| | | Participant | Intercept | 1.36 | | Participant | Intercept | 1.68 |
| | | Residual | | 2.12 | | Residual | | 2.96 |
| | | Observations: 240 | | | | Observations: 240 | | |
| | **lSMA Stimulation** | | | | | | | |
| | **Pre-TMS** | | | | **Post-TMS** | | | |
| **Cued Trials** | **Fixed Effects** | **β** | **SE** | **p($\chi^2$)** | **Fixed Effects** | **β** | **SE** | **p($\chi^2$)** |
| | Intercept | 6.37 | 0.34 | **<0.001** | Intercept | 6.63 | 0.26 | **<0.001** |
| | Serial Order | 0.06 | 0.04 | 0.121 | Serial Order | -0.00 | 0.03 | 0.980 |
| | **Random effects** | **Groups** | | **SD** | **Random effects** | **Groups** | | **SD** |
| | | Participant | Intercept | 1.06 | | Participant | Intercept | 0.75 |
| | | Residual | | 1.86 | | Residual | | 1.53 |
| | | Observations: 240 | | | | Observations: 240 | | |
| **Non-cued Trials** | **Fixed Effects** | **β** | **SE** | **p($\chi^2$)** | **Fixed Effects** | **β** | **SE** | **p($\chi^2$)** |
| | Intercept | 7.64 | 0.46 | **<0.001** | Intercept | 8.06 | 0.46 | **<0.001** |
| | Serial Order | 0.08 | 0.05 | 0.070 | Serial Order | 0.03 | 0.05 | 0.563 |
| | **Random effects** | **Groups** | | **SD** | **Random effects** | **Groups** | | **SD** |
| | | Participant | Intercept | 1.81 | | Participant | Intercept | 1.54 |

| | | | | |
|---|---|---|---|---|
| | Residual<br>Observations:<br>240 | 2.01 | Residual<br>Observations:<br>240 | 2.43 |

**Supplementary Table 8.** Estimated Marginal Means for In-phase (a & c) and Anti-Phase (b & d) Coordination modes.

| **In-Phase Coordination Mode (a) Continuous Relative Phase (ϕ)** | | | | | |
|---|---|---|---|---|---|
| **Pre/Post & Stimulation type** | **Trial type** | **emmean** | **SE** | **lower.CL** | **upper.CL** |
| PreTMS (lSMA) | Cued | 5.40 | 0.18 | 5.04 | 5.77 |
| PostTMS (lSMA) | Cued | 5.33 | 0.18 | 4.97 | 5.70 |
| PreTMS (Sham) | Cued | 5.41 | 0.18 | 5.04 | 5.77 |
| PostTMS (Sham) | Cued | 5.34 | 0.18 | 4.97 | 5.71 |
| PreTMS (lSMA) | Non-cued | 5.63 | 0.18 | 5.26 | 5.99 |
| PostTMS (lSMA) | Non-cued | 5.56 | 0.18 | 5.19 | 5.92 |
| PreTMS (Sham) | Non-cued | 5.56 | 0.18 | 5.20 | 5.93 |
| PostTMS (Sham) | Non-cued | 5.63 | 0.18 | 5.27 | 6.00 |
| Confidence level used: 0.95 | | | | | |

| **Anti-Phase Coordination Mode (b) Continuous Relative Phase (ϕ)** | | | | | |
|---|---|---|---|---|---|
| **Pre/Post & Stimulation type** | **Trial type** | **emmean** | **SE** | **lower.CL** | **upper.CL** |
| PreTMS (lSMA) | Cued | 171.7 | 0.35 | 171.0 | 172.4 |
| PostTMS (lSMA) | Cued | 171.6 | 0.35 | 170.9 | 172.3 |
| PreTMS (Sham) | Cued | 171.7 | 0.35 | 171.0 | 172.4 |
| PostTMS (Sham) | Cued | 171.6 | 0.35 | 170.9 | 172.3 |
| PreTMS (lSMA) | Non-cued | 169.8 | 0.35 | 169.1 | 170.5 |
| PostTMS (lSMA) | Non-cued | 169.7 | 0.35 | 169.0 | 170.4 |
| PreTMS (Sham) | Non-cued | 169.8 | 0.35 | 169.1 | 170.5 |
| PostTMS (Sham) | Non-cued | 169.7 | 0.35 | 169.0 | 170.4 |
| Confidence level used: 0.95 | | | | | |

| **In-Phase Coordination Mode (c) Variability of Continuous Relative Phase (SDϕ)** | | | | | |
|---|---|---|---|---|---|
| **Pre/Post & Stimulation type** | **Trial type** | **emmean** | **SE** | **lower.CL** | **upper.CL** |
| PreTMS (lSMA) | Cued | 4.39 | 0.16 | 4.06 | 4.73 |
| PostTMS (lSMA) | Cued | 4.35 | 0.16 | 4.02 | 4.69 |
| PreTMS (Sham) | Cued | 4.40 | 0.16 | 4.07 | 4.73 |
| PostTMS (Sham) | Cued | 4.36 | 0.16 | 4.03 | 4.69 |
| PreTMS (lSMA) | Non-cued | 4.63 | 0.16 | 4.30 | 4.97 |
| PostTMS (lSMA) | Non-cued | 4.60 | 0.16 | 4.26 | 4.93 |
| PreTMS (Sham) | Non-cued | 4.64 | 0.16 | 4.31 | 4.97 |
| PostTMS (Sham) | Non-cued | 4.60 | 0.16 | 4.27 | 4.93 |
| Confidence level used: 0.95 | | | | | |

| **Anti-Phase Coordination Mode (d) Variability of Continuous Relative Phase (SDϕ)** | | | | | |
|---|---|---|---|---|---|
| **Pre/Post & Stimulation type** | **Trial type** | **emmean** | **SE** | **lower.CL** | **upper.CL** |
| PreTMS (lSMA) | Cued | 6.60 | 0.25 | 6.10 | 7.11 |
| PostTMS (lSMA) | Cued | 6.72 | 0.25 | 6.22 | 7.22 |
| PreTMS (Sham) | Cued | 6.60 | 0.25 | 6.10 | 7.11 |
| PostTMS (Sham) | Cued | 6.72 | 0.25 | 6.22 | 7.23 |
| PreTMS (lSMA) | Non-cued | 8.12 | 0.25 | 7.62 | 8.62 |
| PostTMS (lSMA) | Non-cued | 8.24 | 0.25 | 7.73 | 8.74 |
| PreTMS (Sham) | Non-cued | 8.12 | 0.25 | 7.62 | 8.63 |
| PostTMS (Sham) | Non-cued | 8.24 | 0.25 | 7.73 | 8.74 |
| Confidence level used: 0.95 | | | | | |

### Supplementary Figure 2.

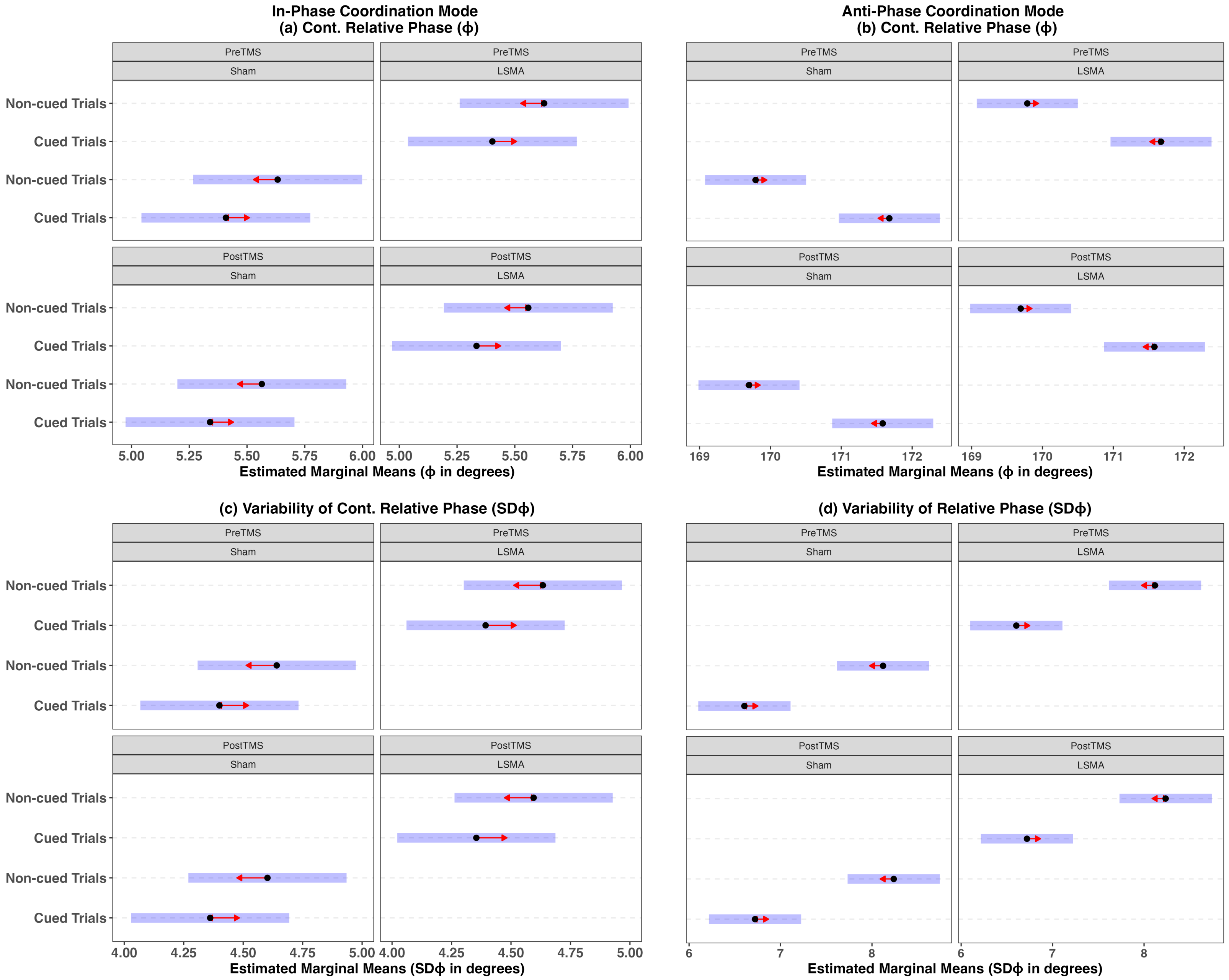

**S2.** Pairwise Comparisons of Estimated Marginal Means for Continuous Relative Phase ($\phi$) and Variability of Continuous Relative Phase (SD$\phi$) in Two Coordination Modes: (a) Continuous Relative Phase in the In-Phase Coordination Mode, (b) Continuous Relative Phase in the Anti-Phase Coordination Mode, (c) Variability in the In-Phase Coordination Mode, and (d) Variability in the Anti-Phase Coordination Mode. The blue bars indicate 95% confidence intervals. Red arrows depict mean comparisons. Non-significant differences are indicated when red arrows overlap with arrows from other categories.
